# Chromatin Compaction Follows a Power Law Scaling with Cell Size from Interphase Through Mitosis

**DOI:** 10.1101/2024.12.05.627016

**Authors:** Petra Stockinger, Anna Oddone, Melike Lakadamyali, Manuel Mendoza, Jérôme Solon

**Affiliations:** Centre for Genomic Regulation (CRG), The Barcelona Institute of Science and Technology, Barcelona, Spain; Universitat Pompeu Fabra (UPF), Barcelona, Spain; ICFO, Institut de Ciènces fotòniques, The Barcelona Institute of Science and Technology, Barcelona, Spain; Department of Physiology, Perelman School of Medicine, University of Pennsylvania (UPenn), Philadelphia, USA; Institut de Génétique et de Biologie Moléculaire et Cellulaire, Illkirch, France; Centre National de la Recherche Scientifique, UMR7104, Illkirch, France; Institut National de la Santé et de la Recherche Médicale, U964, Illkirch, France; Université de Strasbourg, Strasbourg, France; Instituto Biofisika (CSIC, UPV/EHU), Basque Excellence Research Centre, Barrio Sarriena, 48940, Leioa, Spain; Ikerbasque, Basque Foundation for Science, 48013 Bilbao, Spain

## Abstract

Coordination of mitotic chromosome compaction with cell size is crucial for proper genome segregation during mitosis. During development, DNA content remains constant but cell size evolves, necessitating a mechanism that scales chromosome compaction with cell size. In this study, we examined chromatin compaction in the developing *Drosophila* nervous system by analyzing the large neuronal stem cells and their smaller progeny, the ganglion mother cells. Using super-resolution 3D Stochastic Optical Reconstruction Microscopy and quantitative time-lapse fluorescence microscopy, we observed that nanoscale chromatin density during interphase scales with nuclear volume according to a power law. This scaling relationship is disrupted by inhibiting histone deacetylase activity, indicating that molecular cues rather than mechanical constraints primarily regulate chromatin compaction. Notably, this power law dependency is maintained into mitosis but the scaling exponent decreases, suggesting phase separation of chromatin events during mitotic compaction. We propose that the scaling of mitotic chromosome size relative to cell size depends on the power law behaviour of interphase chromatin volume, and that scaling of mitotic chromatin compaction is an emergent property of linear polymers undergoing phase separation with their solvent.

**Statement of Significance:** Understanding how chromatin compaction changes with cell size is essential for understanding the mechanisms ensuring accurate genome segregation during cell division. In this study, we combine Voronoi tessellation analysis of Stochastic Optical Reconstruction Microscopy and image processing methods based on fluorescence intensity spectrum analysis of live confocal images to measure chromatin volume in interphase and mitosis in cells of different sizes. This work reveals that chromatin density in *Drosophila* neuronal cells follows a power law relationship with nuclear volume from interphase through mitosis, suggesting a fundamental principle underlying chromosome scaling across different cell sizes. We show that this scaling is regulated by molecular cues, such as histone deacetylase activity, and is conserved through a potential phase separation during mitosis, providing new insights into the biophysical processes that govern chromatin organization. This work broadens our understanding of chromosome biology and will have implications for understanding size-dependent chromatin dynamics in other organisms.

## Introduction

Proper segregation of the genome into daughter cells during mitosis requires adequate compaction of interphase chromatin into condensed mitotic chromosomes. Since cell size changes during development, but DNA content does not, mitotic chromosome compaction must be coordinated with cell size to ensure proper separation of daughter cell genomes. Indeed, chromosome length scales with cell size in budding yeast, nematodes, and amphibians (Hara et al., 2013; Kieserman & Heald, 2011; Ladouceur et al., 2015, 2017; Neurohr et al., 2011; Zhou et al., 2023), but the scaling mechanisms remain unclear. Here, we investigated chromosome scaling in *Drosophila* developing nervous system, by analyzing chromatin compaction in large neuronal stem cells and their smaller daughter cells, the ganglion mother cells. To measure chromatin density during interphase and mitosis, we used a combination of 3D Stochastic Optical Reconstruction Microscopy and quantitative time-lapse fluorescence microscopy. Our findings reveal that interphase chromatin compaction scales with nuclear volume following a power law. This scaling is impaired when histone deacetylase activity is inhibited, suggesting that chromatin volume scaling is controlled by molecular cues rather than by nuclear envelope-dependent mechanical constraints. Additionally, we found that the power law dependency persists from interphase to metaphase, although the scaling exponent gradually decreases as cells enter mitosis. This reduction in the scaling exponent suggests a phase separation of the chromatin, akin to a linear polymer switching from a good solvent to a poor solvent environment during mitotic compaction. Therefore, we propose that the scaling of mitotic chromosome size relative to cell size depends on the interphase chromatin volume following a power law, and that scaling of mitotic chromatin compaction is an emergent property of linear polymers undergoing phase separation with their solvent.

## RESULTS

### Anaphase chromosome size scales with cell and nuclear size in *Drosophila* nervous system

To investigate how chromosome compaction changes with cell size during mitosis, we used the neuronal precursor cells of the *Drosophila* larval brain as a model system. In this organ, stem cells of the developing nervous system, called neuroblasts (NBs), undergo repeated rounds of asymmetric divisions, generating cells of different sizes. The most abundant type, Type I NBs, divide asymmetrically to generate another NB and a smaller ganglion mother cell (GMC). The GMC subsequently divides to produce two neurons and/or glial cells (Figure **1A**) (Homem & Knoblich, 2012). Neuroblasts and GMCs cover a 4-6-fold size range (Homem et al., 2013), providing an ideal context to investigate whether and how mitotic chromosome size scales with cell size.

**Figure 1:**
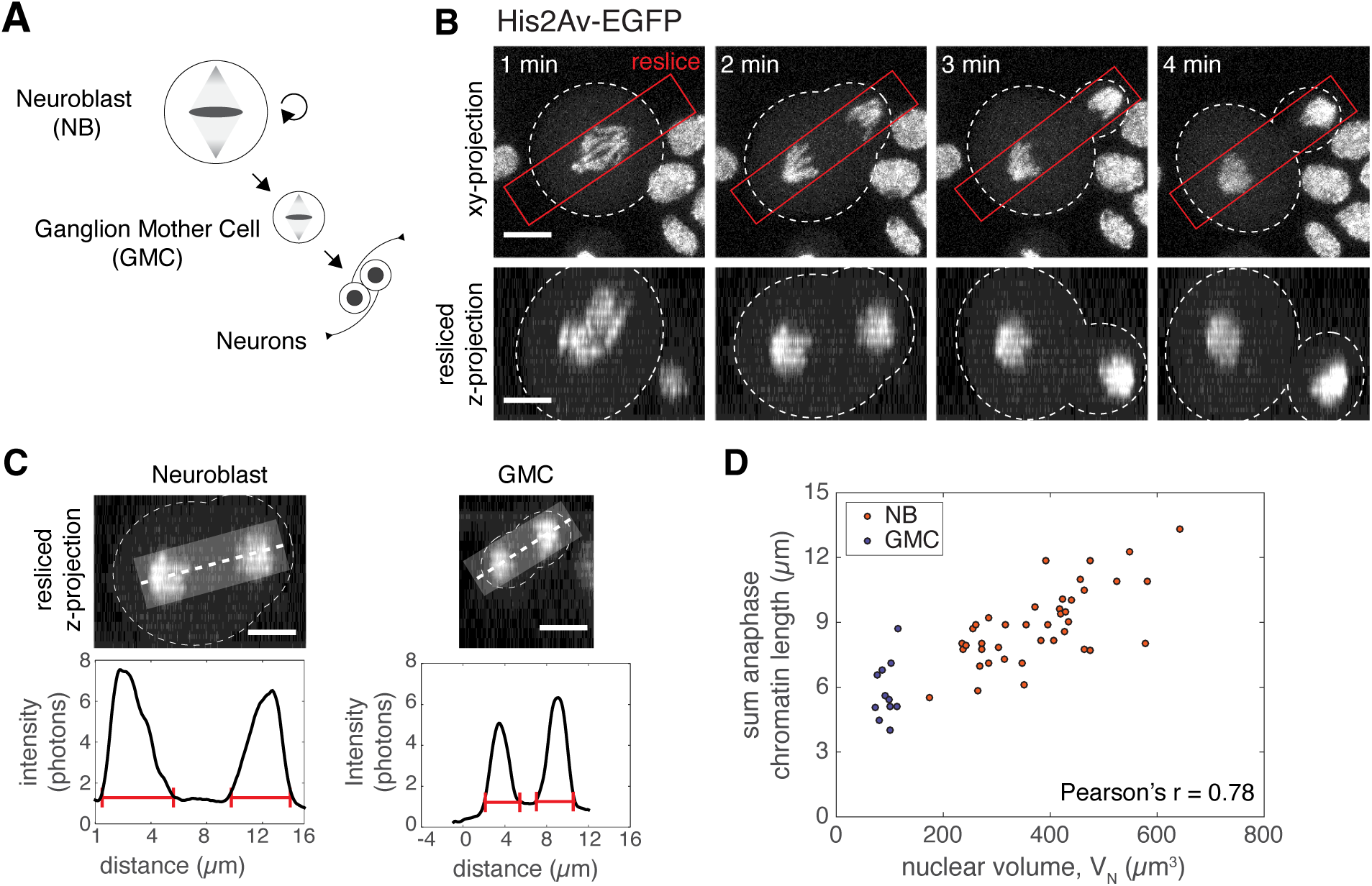
Anaphase chromosome length scales with nuclear volume in the developing D. *melanogaster* nervous system. **A)** Schematic of *Drosophila* type I neuroblast (NB) undergoing asymmetric cell division to form another NB and a ganglion mother cell (GMC). GMCs divide symmetrically to form two neurons/glial cells. **B)** Time series showing the division of chromosomes in an asymmetric division of a neuroblast expressing His2Av-EGFP. The images on the top are mean intensity projections, the dashed line indicates the cell boundaries, and the red rectangle the zone in which reslices have been performed to include the full chromosomal mass. At the bottom, the mean intensity projection of the resliced images along z. Scale bar = 5 μm. **C)** Quantification of anaphase chromatin length. Mean intensity z-projection of a neuroblast and GMC expressing His2Av-EGFP and the respective mean intensity graphs taken along the rectangles centred along the dashed line on the maximum projection images. Total length of the segregating chromatin masses (red segments) was quantified. Scale bar = 5 μm. **D)** Correlation of anaphase chromatin length with nuclear volume (measured 10-20 mins before NEB) in NBs and GMCs. A linear correlation between chromosome length and nuclear volume can be observed. Pearson’s coefficient r=0.77899, p=1.0425588e-11.

To study chromosome compaction in cells of different sizes, we isolated neuronal precursor cells from 3^rd^ instar larval brains expressing GFP-tagged Histone H2A variant (His2Av) proteins. We then acquired 3D time-lapse images using a combination of high-speed confocal imaging and photon-counting techniques to achieve a high signal-to-noise ratio (**Movie 1**, Figure **1B**). The length of the segregating anaphase chromatin mass was used as a proxy for mitotic compaction levels. By re-slicing images along the chromosome segregation axis, we estimated the total length of the segregating chromatin mass in both asymmetrically dividing NBs and in symmetrically dividing GMCs. This allowed us to correlate anaphase chromatin length with spindle length and nuclear volume (see Methods and Figure **1C**).

Anaphase chromatin length ranged from 4 to 13 µm and correlated with interphase nuclear volume (Pearson correlation r=0.78, p=10^−11^) (Figure **1D**). In addition, we observed a correlation between anaphase chromatin length and anaphase spindle length (Pearson correlation r=0.64, p=3×10^−9^) in fixed larval brains (Figure **S1A-B**). Thus, anaphase chromosome size scales linearly with anaphase spindle length, nuclear volume, and cell size in the *Drosophila* central nervous system (CNS), as nuclear size is proportional to cell size in most eukaryotic cells (Conklin, 1912), including *Drosophila* neural precursors (Figure **S1C**). This result suggests that cell size, rather than cell type, is the main determinant of anaphase chromosome size in the Drosophila CNS.

### Scaling of interphase chromatin compaction with nuclear size determined with superresolution microscopy and Voronoi tessellation

Next, we sought to determine whether interphase chromatin compaction scales with cell and nuclear size in *Drosophila* CNS. Previous studies in *C. elegans* and *Xenopus* have shown that mitotic chromosome compaction levels are proportional to nuclear size (Hara et al., 2013; Kieserman & Heald, 2011). However, it is unclear whether nuclear volume affects interphase DNA density. The dense packing of chromatin poses a challenge for conventional optical microscopy, which lacks the resolution needed to investigate genome architecture as it relates to nuclear size. To address this, we used super-resolution microscopy techniques in combination with DNA labeling methods to generate a detailed spatial description of chromatin organization within interphase nuclei (Bintu et al., 2018; Otterstrom et al., 2019; Ricci et al., 2015).

We first labeled DNA with Alexa Fluor 647 (A647) by combining 5-Ethynyl-2′-deoxyuridine (EdU) incorporation and “click’’ chemistry, enabling us to visualize chromatin (Otterstrom et al., 2019; Zessin et al., 2012). To achieve super-resolution in three dimensions, we used Stochastic Optical Reconstruction Microscopy (STORM) in combination with optical astigmatism approaches (Huang et al., 2008). We then analyzed chromatin densities in sections of controlled depth (sections of 50-100 nm, scaled to nuclear size; see Materials and Methods) using Voronoi tessellation (Levet et al., 2015). In this method, a polygon is generated around each localization, where the area of the Voronoi polygon is inversely proportional to the chromatin density in that region (Figure **2A**) (Otterstrom et al., 2019).

**Figure 2:**
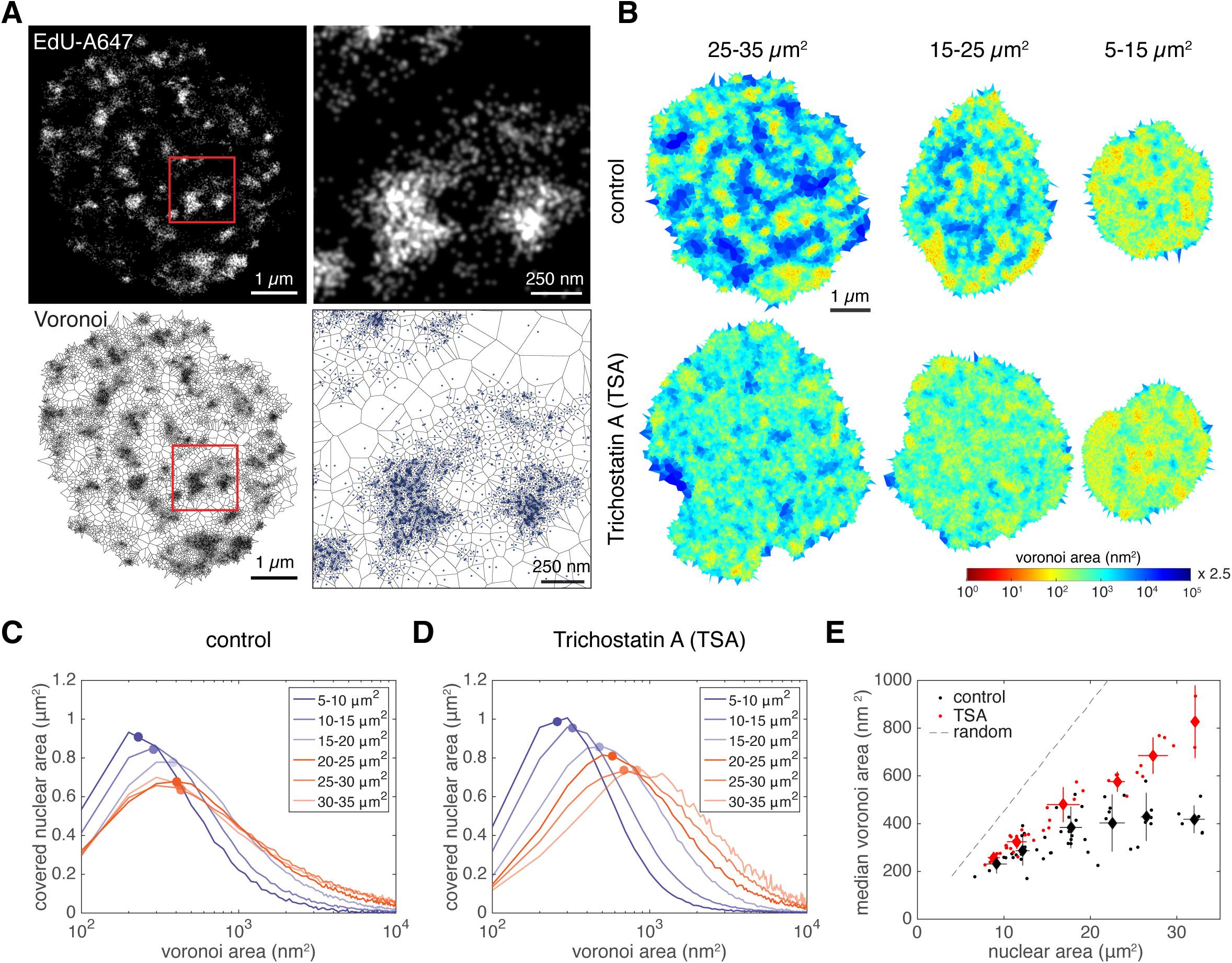
Quantification of interphase chromatin using superresolution microscopy and Voronoi tessellation techniques. **A)** Rendered 3D dSTORM image of a 78 nm section of a neuroblast interphase nucleus labeled with EdU-A647. A higher-zoomed image of the region inside the red square is shown on the right. Lower panels show the data after Voronoi tessellation. **B)** Representative Voronoi diagrams of nuclei of different size ranges in control and TSA-treated cultures. Color coding is according to the size of the Voronoi polygon area. **C, D)** Graphs represent the nuclear area covered by a given Voronoi polygon area in control and TSA-treated cells. Lines represent the average distribution of the binned data (n=2-9 nuclei) and dots indicate the median of the distribution of Voronoi areas of cells of different sizes. **E)** Median Voronoi area of control and TSA treated cell as a function of nuclear area. Diamonds represent the average median polygon area, as well as the median nuclear area ± SD of the binned data in Figure 2 C,D. The dashed line represents the average polygon area of randomly distributed localizations (see Methods).

To compare chromatin densities between neuronal precursor cells (NBs and GMCs) of different sizes, we examined the polygon areas across various nuclear sizes. Large nuclei exhibited significant heterogeneity in Voronoi areas, with high (red), intermediate (orange-green), and low (blue) density regions. In intermediate-sized nuclei, the low-density areas were reduced, and in small nuclei, they were nearly absent (Figure **2B**, “control”). These low-density areas (blue) may correspond to previously described chromatin-free “channels” (Cremer & Cremer, 2001). Since Voronoi areas are inversely proportional to chromatin density, small areas represent high-density regions, whereas large areas represent low-density or chromatin-free regions.

In nuclei smaller than 20 µm^2^, Voronoi areas decreased rapidly with nuclear size, indicating a sharp increase in global chromatin compaction in smaller nuclei (Figure **2B-C**). In contrast, nuclei larger than 20 µm^2^, showed a similar distribution of Voronoi areas, indicating comparable chromatin compaction across these larger nuclei despite differences in nuclear size. We also analyzed the total nuclear area occupied by chromatin (“chromatin-occupied area”) using a simple thresholding method (Figure **S2A**) and observed a similar trend: the chromatin-occupied area scales sub-linearly with nuclear area (Figures **2E** and **S2B**, “control”). Interestingly, when plotting the chromatin-occupied area (A_C_) as a function of the nuclear area (A_N_) on a log-log scale, we observed a linear relationship, indicating that these two variables follow a power law of the form 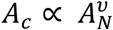 (Figure **S2C**).

Finally, we examined the effects of inhibiting histone deacetylase (HDAC) activity on the power law scaling relationship by treating cells with trichostatin A (TSA), which disrupts interphase chromatin organization (Otterstrom et al., 2019). TSA treatment resulted in a global homogenization of the chromatin signal (Figure **2B**), an expansion of the chromatin-occupied area (Figure **S2B**) and an increase in median Voronoi areas (Figure **2D-E**). Notably, TSA-induced chromatin decompaction occurred without changes to nuclear size (Figure **S2D**). Interestingly, the power law relationship between nuclear and chromatin area observed in untreated cells was lost after TSA treatment, resulting in linear scaling between nuclear size and chromatin density in both small and large cells (Figure **2B** and **S2B**).

In summary, tessellation analysis of STORM images reveals a power law scaling relationship between chromatin volume and nuclear volume during interphase. The effect of TSA treatment suggests that this scaling is driven by internal molecular interactions within the chromatin, rather than by mechanical constraints imposed by the nucleus.

### A method to quantify chromatin compaction dynamics during mitosis

To relate the observed scaling of chromatin and nuclear volumes in interphase to the scaling of mitotic chromosome length according to cell size, we measured chromatin compaction dynamics during mitosis. Accurately measuring chromatin volume in live cells during mitosis poses significant challenges due to the fluctuations in signal intensity caused by chromatin compaction over time. Segmentation-based volume measurement methods are particularly sensitive to threshold values used in the analysis, making them prone to artifacts. To overcome these challenges, we developed a novel method to estimate chromatin volume during confocal live imaging without relying on thresholding. This method uses differences in the intensity distributions of fluorescently labelled histones, distinguishing those that are freely diffusible and those associated with chromatin.

To illustrate our method, we can consider the GFP intensity distribution of His2Av-GFP in cells transitioning from interphase to metaphase (**Figure 3A, B**). This signal captures the contribution of histones bound to the chromatin, as well as free histone molecules that diffuse into the cytoplasm following nuclear envelope breakdown (NEB) (Figure **3A**). To distinguish between signals from chromatin-bound and free histone molecules without manual thresholding, we exploited the different properties of freely diffusible vs. chromosome-associated His2Av-GFP. The intensity distribution of His2Av-GFP fluorescence in the cytoplasm of mitotic cells follows a Poisson probability distribution, consistent with freely diffusing, uncoupled molecules (Figure **3D**). In contrast, the intensity distribution of His2Av-GFP in mitotic chromosomes does not conform to a Poisson distribution and instead requires the addition of a non-Poissonian term, likely due to the tight association of histones with DNA (Figure **3E**). To quantify the contribution of both free and chromatin-bound histone signal, we fitted the His2Av-GFP intensity curves across the whole cell using a two-component distribution function (see Methods, section “Analysis of chromatin volume during mitosis”, and Figure **3C**). Since this method relies on the statistical distribution of different populations of His2Av molecules, it enables the estimation of chromatin volume without the need for thresholding or other arbitrary parameters. To validate this mathematical fit, we independently measured chromatin-bound and free histone populations by manually segmenting chromatin during metaphase, where the cytoplasmic and chromatin-bound histone signals are spatially distinct (Figure **S3A-B**). We observed a strong correlation between the fit and the manual measurements in estimating both the cytoplasmic His2Av concentration and the chromatin-bound His2Av (Figure **S3C-D**). Because the volume assessment performed by our fluorescence intensity spectrum analysis matches manual volume segmentation, we conclude that our intensity distribution-based method allows for determining mitotic chromatin volume in live cells.

**Figure 3:**
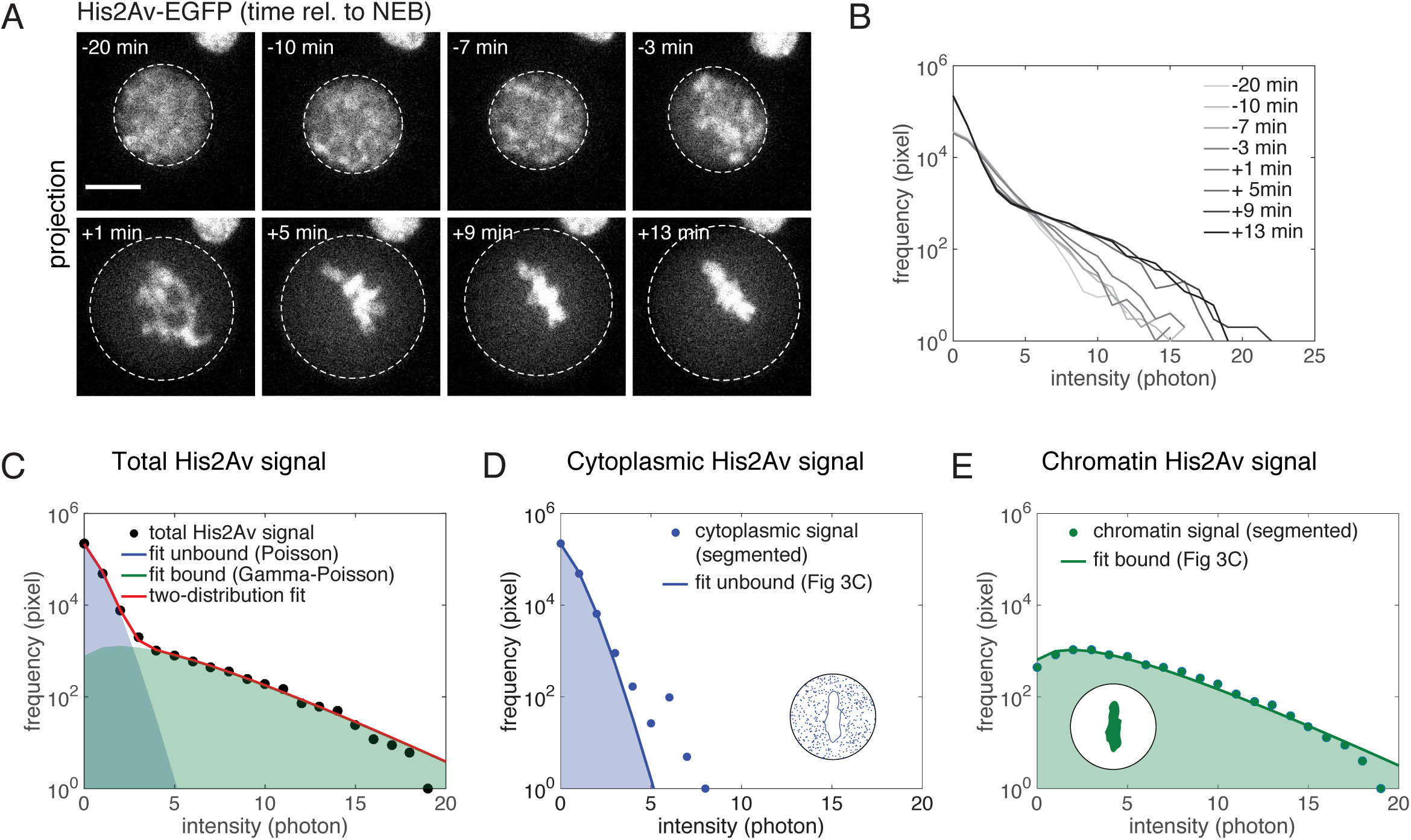
Quantification of chromatin compaction dynamics during mitosis. **A)** Time sequence showing the sum intensity projections of a neuroblast expressing His2Av-EGFP. 0 min indicates the time of nuclear envelope breakdown (NEB). The dashed line highlights the nuclear boundary before NEB and the cell boundary after. Scale bar = 5 µm. **B)** Signal intensity distribution of the nucleus (before NEB) or the entire cell (after NEB) in Figure 3A. **C)** Total signal obtained from the cell in metaphase (+13 min rel. to NEB). Fitting of the total signal with the two-distribution function (red line) as well as the extracted unbound (blue area) and bound (green area) fraction of the His2Av signal. **D)** Cytoplasmic signal extracted by manual segmentation and overlay with the free fraction obtained from the fit in Figure 3C. Measured mean signal intensity = 0.23 photons/pixel (Figure S3A), μ, which is the mean signal from the fit = 0.2592 photons/pixel. **E)** Chromatin signal of the entire nucleus extracted by manual segmentation and overlay with the bound fraction obtained from the fit in Figure 3C.

To be able to correlate chromatin compaction behaviours in interphase with mitosis using different techniques, we estimated chromatin volume from fixed and live samples at similar stages (interphase/prophase). Using the live chromatin compaction method described above, we extracted the average chromatin density, and from this, calculated the chromatin volume during entry into mitosis from time-lapse images (see Methods). We then compared the chromatin volumes during prophase, determined by live imaging, with the chromatin-occupied areas during interphase, determined by STORM imaging (compare Figures **2E**, **S2B**, and **4B**). To estimate chromatin volume from single plane interphase STORM images, we first assumed a spherical geometry of the nucleus. Second, we corrected for the geometrical changes associated with the cell fixation (See Methods, Figure **S4A**). The interphase chromatin volumes estimated by STORM closely matched those obtained from live confocal microscopy, and their dependence on nuclear volume followed the same relationship, validating the accuracy of our methodology (Figure **4A** and Figure **S4B**).

**Figure 4:**
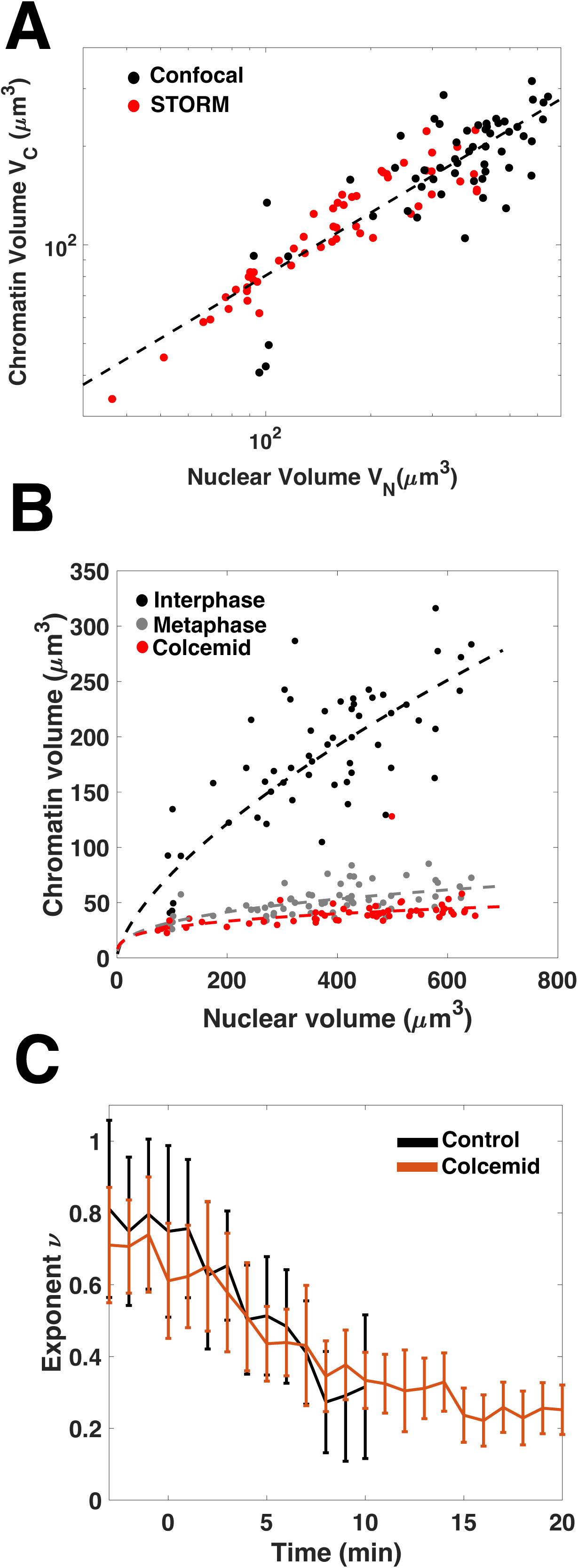
**A)** Graph showing the chromatin volume as a function of the nuclear volume in a log-log scale for fixed STORM data in interphase and live imaging. The two sets of data are in good agreement and follow a power law. A power law fit on the combined dataset is also displayed with the following relation, 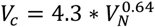 **B)** Graph showing the chromatin volume as a function of the nuclear volume for cells of different sizes obtained with live confocal imaging. The graph shows a power law dependency of the chromatin volume at three different time points, interphase, metaphase, and in extended pro-metaphase in colcemid-treated cells. The respective power law fits are displayed with the following functions: 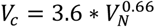 for interphase, 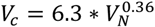 for metaphase and 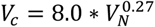 for colcemid-treated cells. **C)** Graph showing the change in fitting power-law exponent over time during mitosis (after NEB) for cells of different sizes in control and colcemid-treated cells. We observe a reduction in both cases from approximately 0.7 before NEB to 0.3 at metaphase.

### The scaling of chromatin volume with nuclear size is maintained during mitosis and follows a power law with a decreasing exponent

Our results demonstrate that chromatin volume in interphase scales with nuclear volume in a similar manner when assessed by both STORM and live cell imaging. This relationship follows a power law with an exponent of approximately 2/3 (exponent value is 0.6674 for STORM and 0.6627 for live imaging; **Figures 4A, 4B** (“interphase”) and Figure **S4C-D**). Similarly, the chromatin volume during metaphase also scales with nuclear volume, but with a lower exponent of approximately ⅓ (exponent value is 0.3572; Figure **4B**, “metaphase”).

Interestingly, a comparable power law dependency is observed in polymer volume as a function of its length, with different possible exponents (see next section) (De Gennes, 1979). To explore whether an additional reduction in the exponent occurs if metaphase is prolonged, we treated cells with the microtubule polymerisation inhibitor colcemid, which blocks anaphase onset (Rieder & Palazzo, 1992). In these cells, compaction quantitatively matched that of untreated cells and persisted for up to 20 min after NEB. We observed a power law relationship in these cells with a further reduced exponent (Figure **4B**, “colcemid”).

Using time-lapse data, we tracked the evolution of the power law exponent at various time points during mitosis. This analysis revealed a continuous decrease in the exponent from prophase to metaphase in both untreated and colcemid-treated cells (Figure **4C**). Thus, the scaling of chromatin volume relative to nuclear volume can be described by power laws with different exponents: approximately 2/3 during interphase, decreasing to 1/3 during metaphase, and continuing to decrease when metaphase is extended.

Together, these results indicate that in *Drosophila* neural progenitor cells, the scaling of mitotic chromosome volume relative to nuclear volume is established during interphase following a power law, and that mitotic compaction preserves this power law dependency with a linear reduction in the exponent over time.

### Mitotic chromatin compaction is consistent with a phase separation of a polymer with its solvent

The power law dependency of chromatin volume as a function of nuclear volume can be interpreted through the lens of polymer physics. Super-resolution imaging of *Drosophila* cells has shown that the length of chromatin domains scales with their volume according to a power law, indicating that chromatin can be effectively modelled using polymer models (Boettiger et al., 2016). The volume occupied by a free polymer is determined by its radius of gyration (R_G_) which is proportional to the polymer’s length *(L)* and follows the power law *R_G_* ∝ *L^V^*, where the exponent *v* depends on the polymer’s solubility. For a polymer in a good solvent, 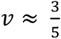, and for a poor solvent 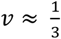 (De Gennes, 1979).

Our measurements suggest that chromatin is not strictly confined by the nucleus, as indicated by TSA-induced decompaction, which affects the power law dependency (see Figure **S2B**). Therefore, the observed power law dependency as a function of nuclear volume appears to emerge from the internal organization of the chromatin itself.

The observed relationship in Figure **4A**, which follows 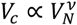, with ν the power law exponent that varies from 2/3 to 1/3, can be rewritten in terms of gyration radii as 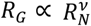, where *R_G_* is the gyration radius of chromatin and *R_N_* is the nuclear radius. This mirrors the laws governing polymer occupancy, suggesting that the effective length of chromatin, as defined by its gyration radius, is proportional to the nuclear radius. In other words, the extent of chromatin compaction in interphase, which influences its effective length, is dependent on the nuclear radius.

During mitosis, we observe a change in the power law exponent from approximately 0.6 to 0.3. This shift aligns with a phase separation process, where chromatin transitions from a state resembling a good solvent to one resembling a bad solvent (De Gennes, 1979). This is further supported by the emergence of a bimodal distribution of fluorescence intensities for free and bound histones during mitosis, indicating a spatial separation (demixing) of these two forms of the protein (see Figure **3B**).

## DISCUSSION

In summary, our work identifies that chromatin volume scale with cell size following a power law and that the transition from interphase to mitosis consists in a reduction of the power law exponent. Our data suggest that the scaling of mitotic chromatin compaction with cell size is established during interphase. This observation is consistent with recent studies on Xenopus chromosomes, which demonstrated that mitotic chromosomes scale with the nuclear-cytoplasmic ratio and cell size during development (Zhou et al., 2023).

Our findings also suggest that chromosome scaling is an emergent property of linear polymers undergoing phase separation with their solvent. In this model, interphase chromatin structure scales with nuclear volume according to a power law, where the exponent depends on the biochemical environment present during interphase.

How does nuclear volume influence chromatin compaction during interphase? We hypothesize that a smaller chromatin volume increases the likelihood of a DNA chain encountering a compaction factor at distant sites. This concept aligns with existing models of mitotic chromosome condensation, such as the “diffusion capture” model. In this model, compaction factors like condensin complexes stabilize stochastic pairwise interactions between condensin binding sites, contributing to mitotic chromosome formation (Cheng et al., 2015; Gerguri et al., 2021). Similarly, these compaction factors may influence chromatin volume during interphase. As cells transition into mitosis, changes in the biochemical properties of the cell lead to chromatin dewetting and phase separation, resulting in mitotic chromatin compaction scaling to nuclear volume with the same power law as in interphase but with a different exponent.

How is this reduction in the power law exponent during mitosis achieved on a molecular level? Several changes in chromatin properties and organization occur during mitosis, including histone modifications, condensin activity, and surfactant localization. Future studies will need to explore the relative impact of these factors on the reduction in the exponent associated with chromatin phase separation.

## Acknowledgements

We thank Nacho Molina, Anne-Cecile Reymann and Ion Andreu for critical reading of the manuscript, and the CRG microscopy core facility.

This study was supported by the European Research Council (ERC) Starting Grant 2010-St-20091118 to M.M., the European Research Council (ERC) Starting Grant 337191-MOTOS To M.L., and the Spanish Ministry of Economy and Competitiveness, ‘Centro de Excelencia Severo Ochoa 2013–2017’, SEV-2012-0208 to the CRG. Work in the Mendoza lab is supported by ANR-22-CE12-0021 and by Fondation ARC PGA2022010004436_4876. IGBMC, as part of the Interdisciplinary Thematic Institute IMCBio+ 2021-2028 program of the University of Strasbourg, CNRS and Inserm, is also supported by IdEx Unistra (ANR-10-IDEX-0002), SFRI-STRAT’US project (ANR 20-SFRI-0012) and EUR IMCBio (ANR-17-EURE-0023) under the framework of the France 2030 Program. Work in the Solon lab is supported by the Spanish Ministry of Science and Innovation PID2019-109117GB-100 and PID2022-142779NB-I00 and by the Basque government (PIBA_2023_1_0033).

## Materials and Methods

### KEY RESOURCES TABLE

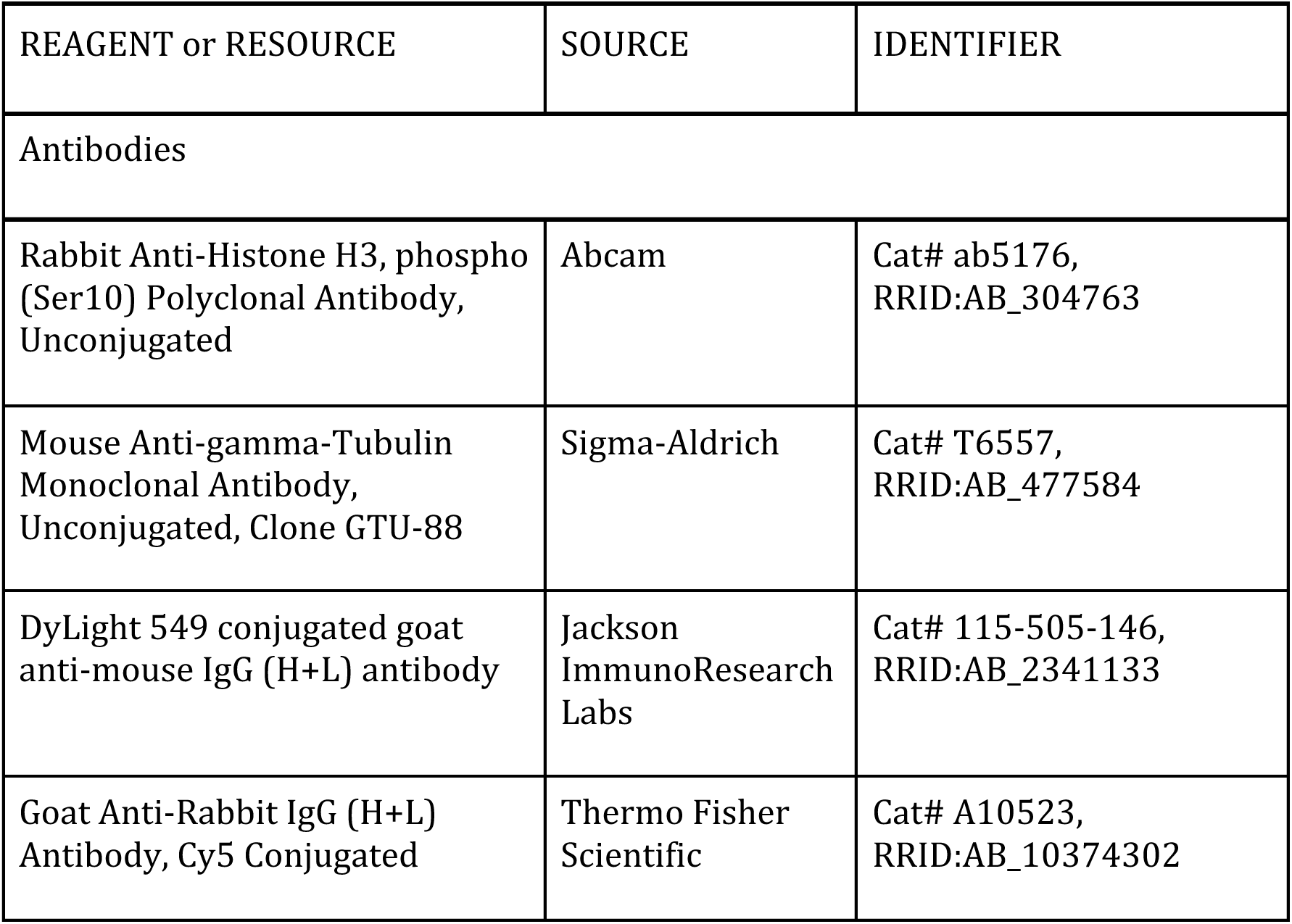
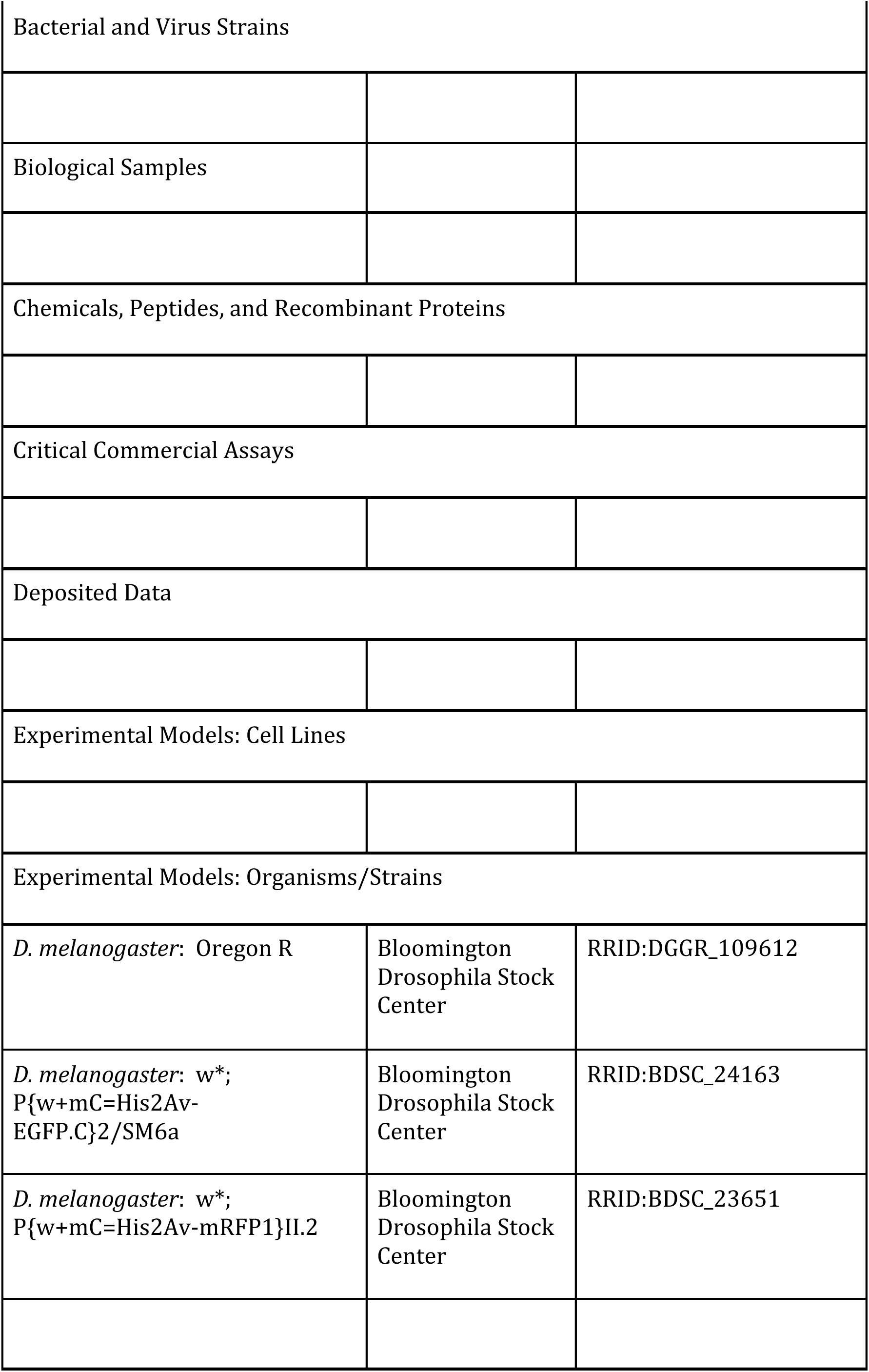
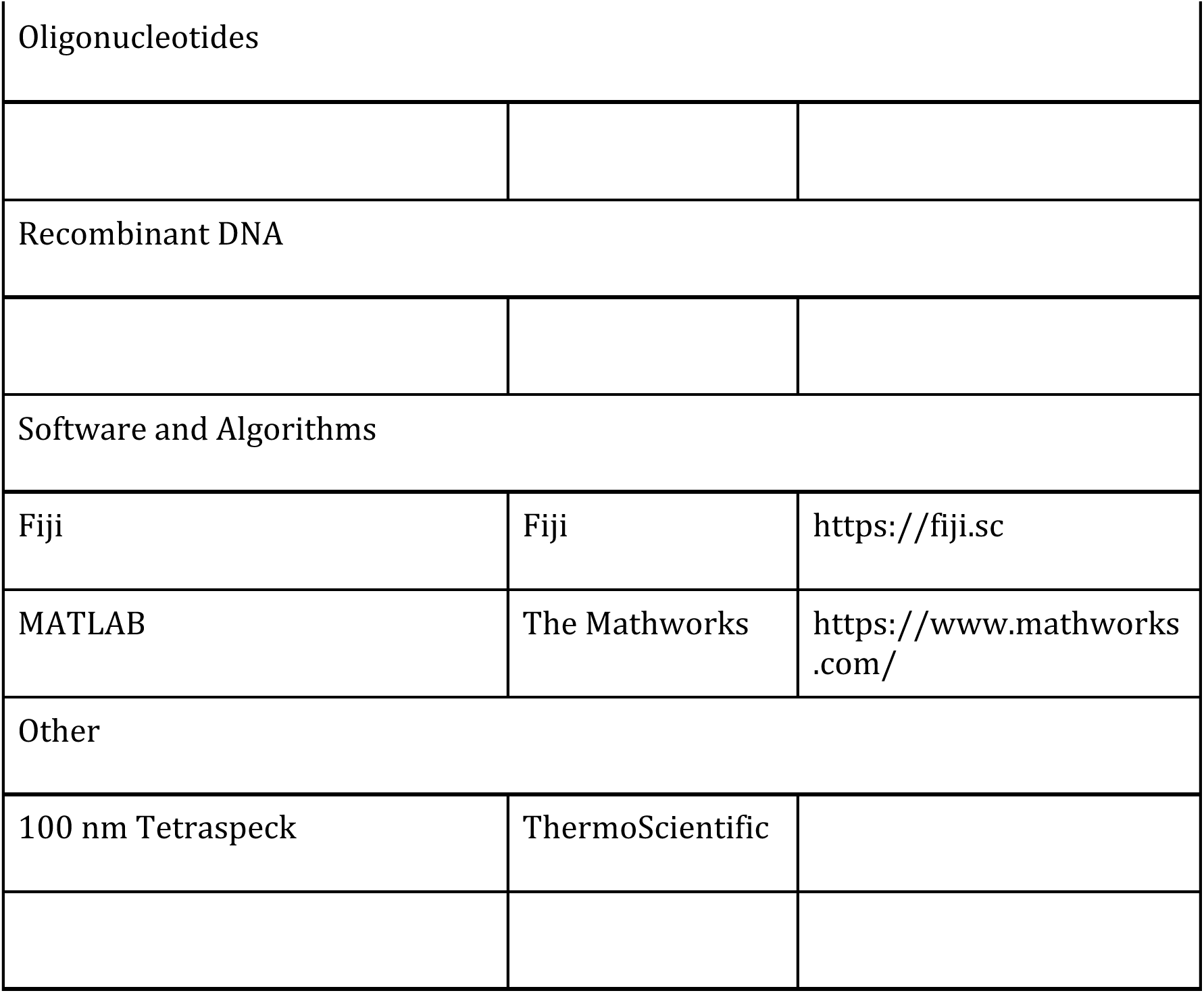

| REAGENT or RESOURCE | SOURCE | IDENTIFIER |
| --- | --- | --- |
| Antibodies |  |  |
| Rabbit Anti-Histone H3, phospho (Ser10) Polyclonal Antibody, Unconjugated | Abcam | Cat# ab5176, RRID:AB_304763 |
| Mouse Anti-gamma-Tubulin Monoclonal Antibody, Unconjugated, Clone GTU-88 | Sigma-Aldrich | Cat# T6557, RRID:AB_477584 |
| DyLight 549 conjugated goat anti-mouse IgG (H+L) antibody | Jackson ImmunoResearch Labs | Cat# 115-505-146, RRID:AB_2341133 |
| Goat Anti-Rabbit IgG (H+L) Antibody, Cy5 Conjugated | Thermo Fisher Scientific | Cat# A10523, RRID:AB_10374302 |
| Bacterial and Virus Strains |  |  |
| Biological Samples |  |  |
| Chemicals, Peptides, and Recombinant Proteins |  |  |
| Critical Commercial Assays |  |  |
| Deposited Data |  |  |
| Experimental Models: Cell Lines |  |  |
| Experimental Models: Organisms/Strains |  |  |
| *D. melanogaster*: Oregon R | Bloomington Drosophila Stock Center | RRID:DGGR_109612 |
| *D. melanogaster*: w*;<br>P{w+mC=His2Av-EGFP.C}2/SM6a | Bloomington Drosophila Stock Center | RRID:BDSC_24163 |
| *D. melanogaster*: w*;<br>P{w+mC=His2Av-mRFP1}II.2 | Bloomington Drosophila Stock Center | RRID:BDSC_23651 |
| Oligonucleotides |  |  |
| Recombinant DNA |  |  |
| Software and Algorithms |  |  |
| Fiji | Fiji | https://fiji.sc |
| MATLAB | The Mathworks | https://www.mathworks.com/ |
| Other |  |  |
| 100 nm Tetraspeck | ThermoScientific |  |

### CONTACT FOR REAGENT AND RESOURCE SHARING

Further information and requests for reagents may be directed to, and will be fulfilled by the Lead Contact, Jerome Solon

### EXPERIMENTAL MODEL AND SUBJECT DETAILS

#### Drosophila Strains

Following lines were used: Oregon R (Bloomington 109612), w*; P{w+mC=His2Av-EGFP.C}2/SM6a (Bloomington 24163), w*; P{w+mC=His2Av-mRFP1}II.2 (Bloomington 23651).

### METHOD DETAILS

#### Fly strains and husbandry

All fly stocks were maintained by standard methods at room temperature and were grown on a standard fly food. 3^rd^ instar larvae of His2Av-eGFP or His2Av-mRFP expressing flies were used for live-imaging experiments and Oregon R for immunofluorescence and dSTORM experiments. All samples in this study were prepared from male and female larvae.

#### Neuroblast primary cell culture

Larval brains were dissected in cold collagenase buffer (800 mg NaCl, 20 mg KCl, and 5 mg NaH_2_PO_4_, 100 mg NaHCO_3_ and 100 mg Glucose in 100 ml ddH_2_O) and incubated in 0.2 mg/ml collagenase (Sigma) for 20 min at room temperature(Feiguin et al., 1998). Brains were rinsed in collagenase buffer and manually dissociated in Schneider’s medium complemented with 5% FCS, glucose (1mg/ml), 5% fly serum and insulin (5 mg/ml) by pipetting using coated yellow tips (Sigmacote, Sigma).

For live imaging experiments, cells were plated on poly-L-lysine-coated 35 mm glass bottom Fluorodish culture plates (WPI) and allowed to settle for 30 mins at room temperature. Drugs were dissolved in supplemented Schneider’s medium and added to the samples 30 mins before imaging to reach a final concentration of (0.5 μM TSA and/or 1 μM colcemid).

For STORM imaging, cells were plated on poly-L-lysine (Sigma) coated 8-well Chambered Coverglass (Nunc^TM^ Labtek^TM^) and allowed to settle for 30 mins at room temperature before fixation. For drug treatments, the drugs were dissolved in supplemented Schneider’s medium and were added for 2 hours (DMSO, 0.5 or 5 μM TSA) before fixation.

#### Immunofluorescence of larval brains

Larval brains were dissected in cold PBS and fixed with 4% PFA in 0.3% Triton in PBS (PBST) for 20 mins at room temperature. Samples were washed 3x 20min in PBST and blocked in 5% NGS (v/v) in PBST for 1 hour before incubation with primary antibody in blocking solution (overnight, 4°C): rabbit anti-Histone H3 (phospho S10) (1/200); mouse anti-g-tubulin (GTU-88) (1/500).

Brains were washed 3X 10min in PBST and incubated for 2 hours at room temperature with secondary antibody in blocking solution: anti-rabbit Cy5 (1/500); anti-mouse DyLight 549 (1/500). Samples were washed in PBST and incubated for 30min in 50:50 PBS/Glycerol and brains were dissected and mounted in Vectashield (VectorLabs).

#### EdU labeling

Labeling of EdU with an azide derivative of Alexa Fluor 647 was performed using the Click-iT Plus EdU Alexa Fluor 647 Imaging Kit (C10640, Invitrogen). Young (feeding) L3 larvae were collected and fed overnight with standard flyfood containing EdU (5-ethynyl-2’-deoxyuridine) (Invitrogen) and Bromophenol Blue sodium salt (Sigma). Brains of wandering 3^rd^ instar larvae with blue coloured guts were collected and primary brain cultures were prepared as described previously. For imaging experiments 100-200 ml culture (~4 brains) were plated on poly-L-lysine (Sigma) coated 8-well Chambered Coverglass (Nunc^TM^ Labtek^TM^) dishes.

Cells were fixed for 10 min with 4% PFA in PBS and washed twice with 3% BSA in PBS. Fixed cells were rinsed with 1mM glycine and incubated for 10 min in 0.1N HCl. Cells were then washed and permeabilized with 0.3% Triton X-100 in PBS (PBST) for 20 min at room temperature. Samples were washed with 3% BSA in PBS and incubated in 200 ml of freshly prepared Click-iT Plus reaction for 30 min at room temperature. Cells were washed with 3% BSA in PBS and 3x 20 mins with PBST and stored in PBS.

#### Microscopy

##### Confocal microscopy

Live imaging of primary brain cultures was performed at room temperature (22-25°C) using a Leica TCS SP5 II CW-STED (inverted) confocal microscope with a 63x/NA1.4 oil objective using a 488 Argon laser.

To allow high-speed as well as quantitative imaging, a combination of resonant scanning (8000 Hz) with super-sensitive Hybrid detectors (HyD) in photon counting mode were used. Images of 512×512 pixels were acquired with 8x accumulation and 80 nm pixel size. Cells were imaged with 15-20 z-sections at 1 μm distance and 60 s intervals for up to 4 hours.

Imaging of fixed and immunostained larval brains was performed on a Leica TCS SP5 AOBS inverted scanning confocal microscope, using a 63x/NA1.4 Oil objective. Z stacks were acquired with 80 nm pixel size with 251.7 nm z-steps using a 561 nm and a 633 nm laser.

#### 3D dSTORM imaging

Imaging experiments were carried out with a commercial STORM microscope system from Nikon Instruments (N-STORM) equipped with a cylindrical lens for 3D-imaging (Huang *et al*., 2008). Imaging was performed using a standard STORM imaging buffer containing 100 mM Cysteamine MEA, 0.5 mg/mL glucose oxidase, 40 μg/mL catalase, and 5% Glucose (all Sigma-Aldrich) in PBS. An inclined illumination mode was used to obtain the images (Tokunaga et al., 2008). Laser light at 647 nm was used for exciting Alexa Fluor 647, and laser light at 405 nm was used for reactivating it. The emitted light was collected by an oil immersion objective (Apo internal reflection fluorescence 100×; 1.49 N.A.) filtered by an emission filter (ET705/72m) and imaged onto an EMCCD camera. STORM images were analyzed using custom-written software (Insight3; provided by Bo Huang, University of California, San Francisco, CA) by fitting the PSF of individual fluorophores with a simple Gaussian curve in every frame to determine the x and y coordinates.

For 3D-STORM imaging, calibration data were acquired by imaging subdiffractional limit size beads (100 nm Tetraspeck, ThermoScientific) in PBS adsorbed to 8-well Labtek chambers (Nunc) at low enough dilution to enable single bead visualization, using the NIS software STORM module.

### QUANTIFICATION AND STATISTICAL ANALYSIS

The statistical details of experiments are described in the figure legends.

#### Image Analysis

##### Anaphase chromatin length measurements

To measure chromosome size during anaphase in live cells as well as fixed tissues, we analyzed the time point where segregating chromatin masses were fully separated. First, the z-stacks were resliced with 80 nm spacing along the axis of chromosome segregation using the Fiji ‘Reslice’ function and maximum z-projections of the resliced images were obtained. To measure the length of chromatin, a line that covers the width of the chromatin (see Figure 1B), was drawn along the segregation axis. The intensity distribution was extracted by using the ‘Plot Profile’ function and the sum of the chromatin length was measured using a semi-automated script developed in Python.

##### Analysis of chromatin densities in 3D STORM images

First, we manually segmented the nucleus in the rendered images using Fiji and then extracted the localizations within the ROI.

Since the axial localization precision degraded rapidly with distance to the focal plane, we were not able to directly analyze the whole range of 3D data. Therefore, we only analyzed the localization within the focal plane. To compare chromatin densities of nuclei with different sizes, we first scaled the analyzed nuclear volumes to the nuclear size, where the height of the individual volumes is proportional to the nuclear area (h ~ 50-100nm). In addition, we only analyzed 20.000 localisations/volume to directly compare the Voronoi areas between the different nuclei. The resulting single-molecule localizations of each slice were converted into a 2D Voronoi diagram in Matlab. Voronoi diagrams were constructed for each xy coordinate using the ‘Voronoi’ function and Voronoi polygon areas were determined using the ‘polyarea’ function.

To estimate the chromatin-occupied area within individual nuclei, we thresholded the dataset at 10^4^ nm^2^. The histograms of the covered nuclear area were obtained by multiplying the density distribution with the given Voronoi area and binning the data into 100 nm^2^ bins. For each data point (nucleus), we averaged the data of the central 3-5 data sets (“slices”).

##### Analysis of chromatin volume during mitosis

The 3D stacks at each time-point were manually segmented and the histograms were extracted using Fiji software. To separate the chromatin-bound from the free histone fraction we took advantage of the difference in the statistical distribution of the signal due to the different physical behaviors of chromatin-bound and free histone fractions.

The free histone fraction diffuses in the cytoplasm without spatial correlations between the individual histone molecules, therefore the spatial distribution of the signal captured in the different pixels of the image should follow a Poisson distribution.

We could therefore fit the normalized distribution of intensities in the cytoplasm with the following Poisson distribution for continuous distribution of points:

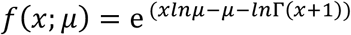

Where the parameter *μ* is the mean of the distribution and the variable x describes the fluorescence intensities (Figure 3C). The fact that the intensity probability distribution of the cytoplasmic histones is well fitted by a Poisson distribution indicates that cytoplasmic histones localizations are spatially uncorrelated (or independent) at the scale of individual pixel and histones can be assumed randomly distributed.

On the other hand, the chromatin bound signal cannot be fitted by a simple Poisson distribution, but can be represented by a Poisson-Gamma mixture,

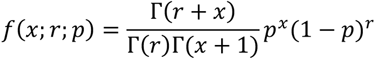

which is a Poisson distribution where the Poisson parameter is itself a random variable, distributed according to a Gamma distribution. The Gamma distribution is described by two parameters: r (defined as the shape of the distribution) and p (defined as the scale of the distribution). The Gamma distribution represents the differences of densities within the chromatin.

Therefore, we can extract the contribution of the free as well as the chromatin bound histone by fitting the intensity curves with a two-distribution function:

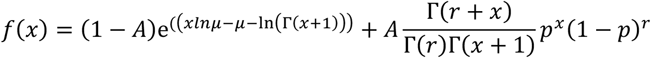

where A is the proportion of pixels occupied by the chromatin.

From the fitting parameters *μ*, *p*, *r*, and *A*, we calculated N, the total signal of bound histone

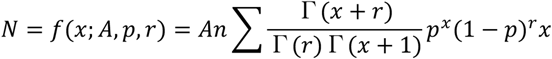

where represents the total number of collected signal.

To quantify the average intensity of the bound histone signal we calculated the mean of the Gamma function

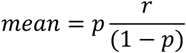

which represents an average histone concentration: mean= N/ V.

Therefore, the average chromatin volume can be calculated from the mean of the Gamma PDF and the total number of bound signal N by

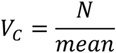

First, fitting the total intensity distribution with our “fitting function” allows us to separate the fraction of the unbound signal from the chromatin bound signal. Second, we can extract the distribution of the chromatin bound signal represented by the Gamma distribution where the mean of the Gamma PDF is similar to the measured mean of the chromatin signal and the number of photons extracted from the fitting are similar to the measured values (Figure S3C).

### DATA AND SOFTWARE AVAILABILITY

Data and software will be provided on demand via email to the corresponding author.

---

**Figure S1:**
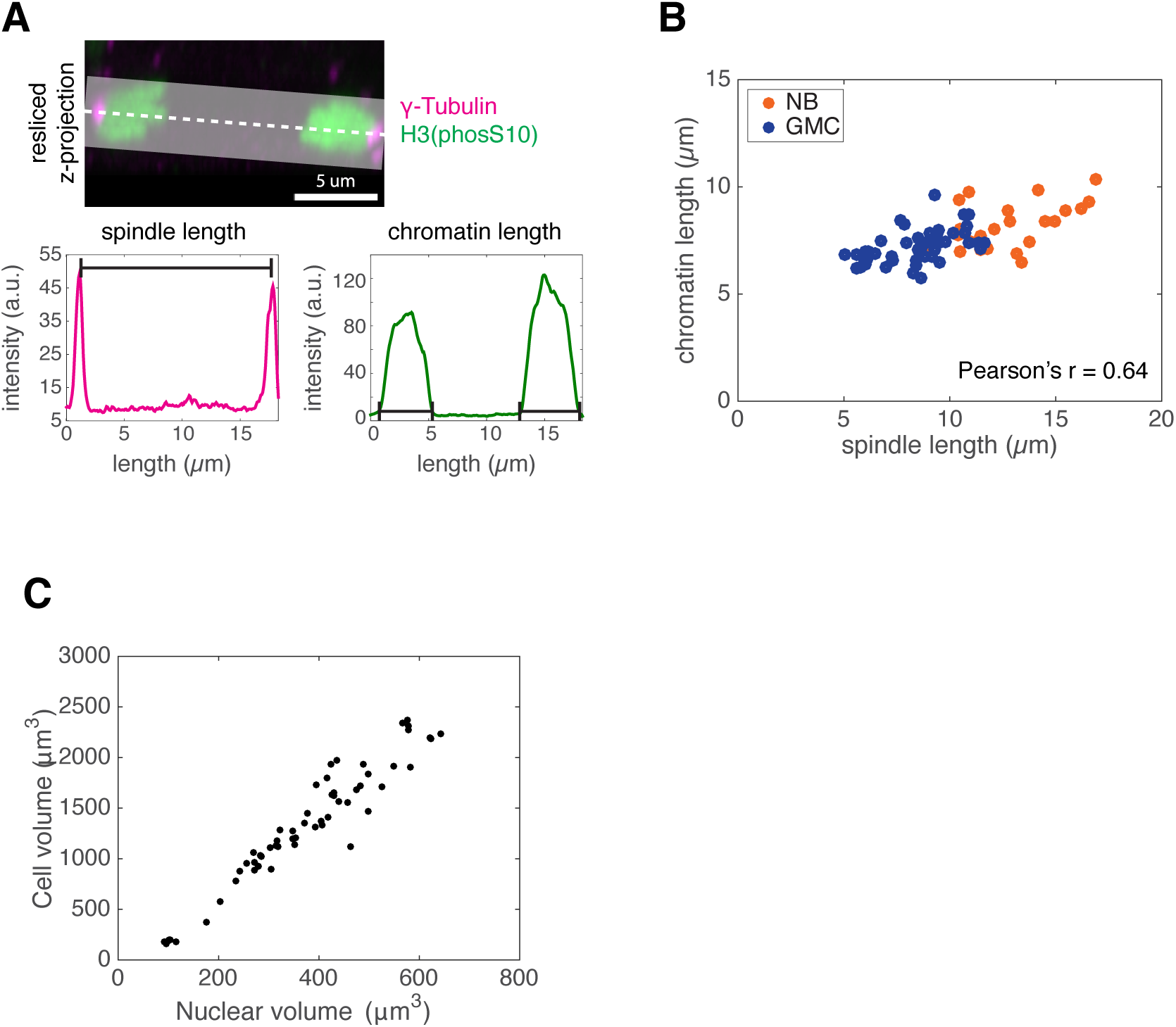
Anaphase chromosome length measurements. **A)** Quantification of spindle length and anaphase chromatin in a neuroblast stained with y-Tubulin (magenta) and anti-Histone H3 (phosS10) (green) antibodies. Mean intensity was measured within the white box. Scale bar = 5 μm. **B)** Correlation of anaphase chromatin length with spindle length (centrosome distance) in NB and GMCs. A linear correlation can be observed. Pearson’s coefficient r=0.636758, p=3.115789e-09. **C)** Graph showing the nuclear volume as a function of the cell volume for NB and GMCs. A linear correlation can be observed between the nuclear and cell volume.

**Figure S2:**
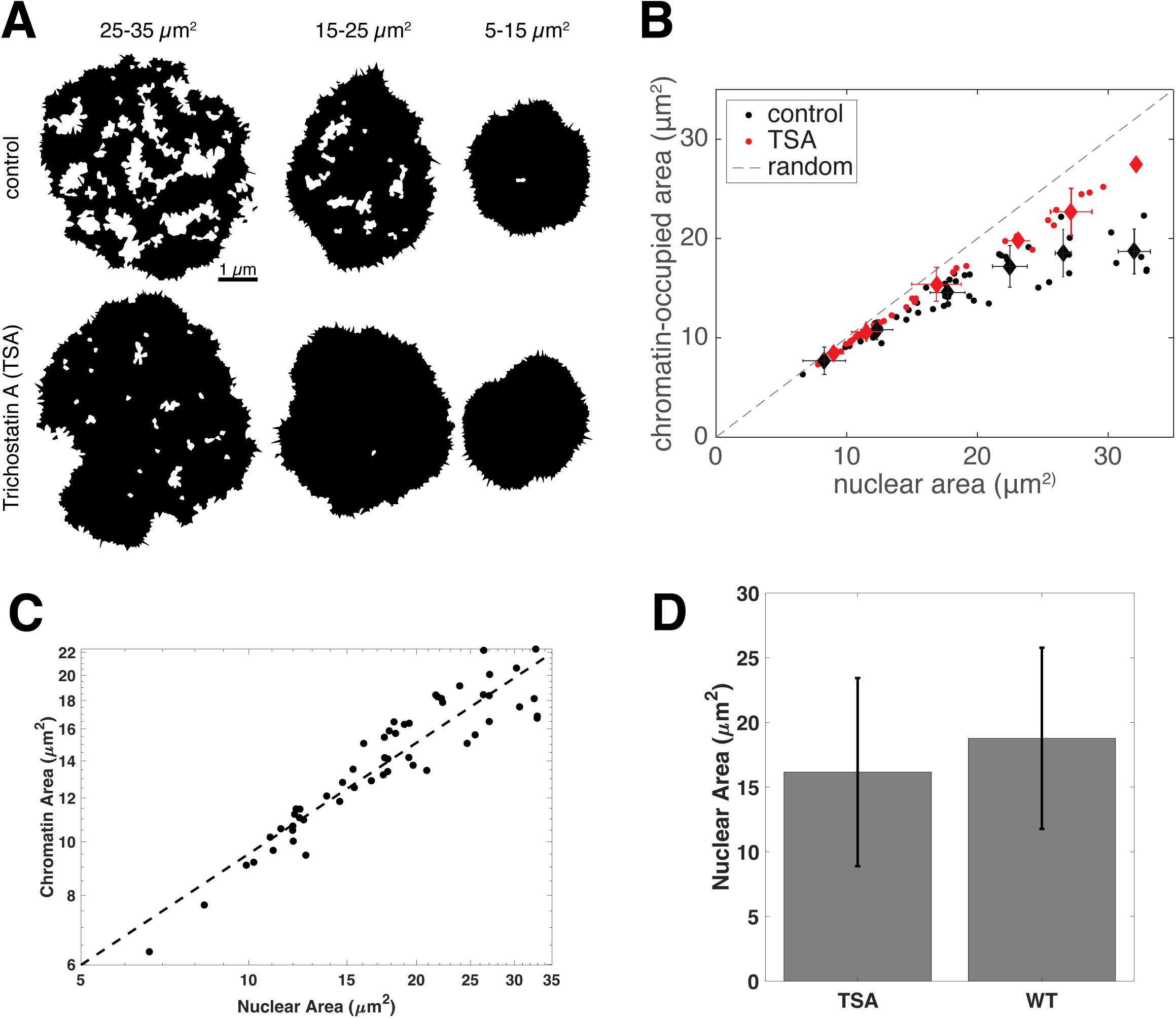
Chromatin occupancy upon thresholding. **A)** Thresholded chromatin-occupied area of the nuclei showed in Figure 2B, after removing polygon areas <10^4^ nm^2^. **B)** Chromatin-occupied area in control and TSA-treated cells. Diamonds represent the average chromatin-occupied area vs nuclear area ± SD of the binned data in Figure S2A. The dashed line represents the area of randomly distributed localizations. **C)** Chromatin-occupied area in control cells as a function of the nuclear area displayed in a log-log plot and corresponding linear plot (dashed line). The dependency of chromatin area with nuclear area follows a power law and can be fitted by the expression, 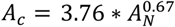 **D)** Nuclear area for control and TSA treated cell. Bars are standard deviations. *N_WT_* = 54 cells; *N_TSA_* = 46 cells.

**Figure S3:**
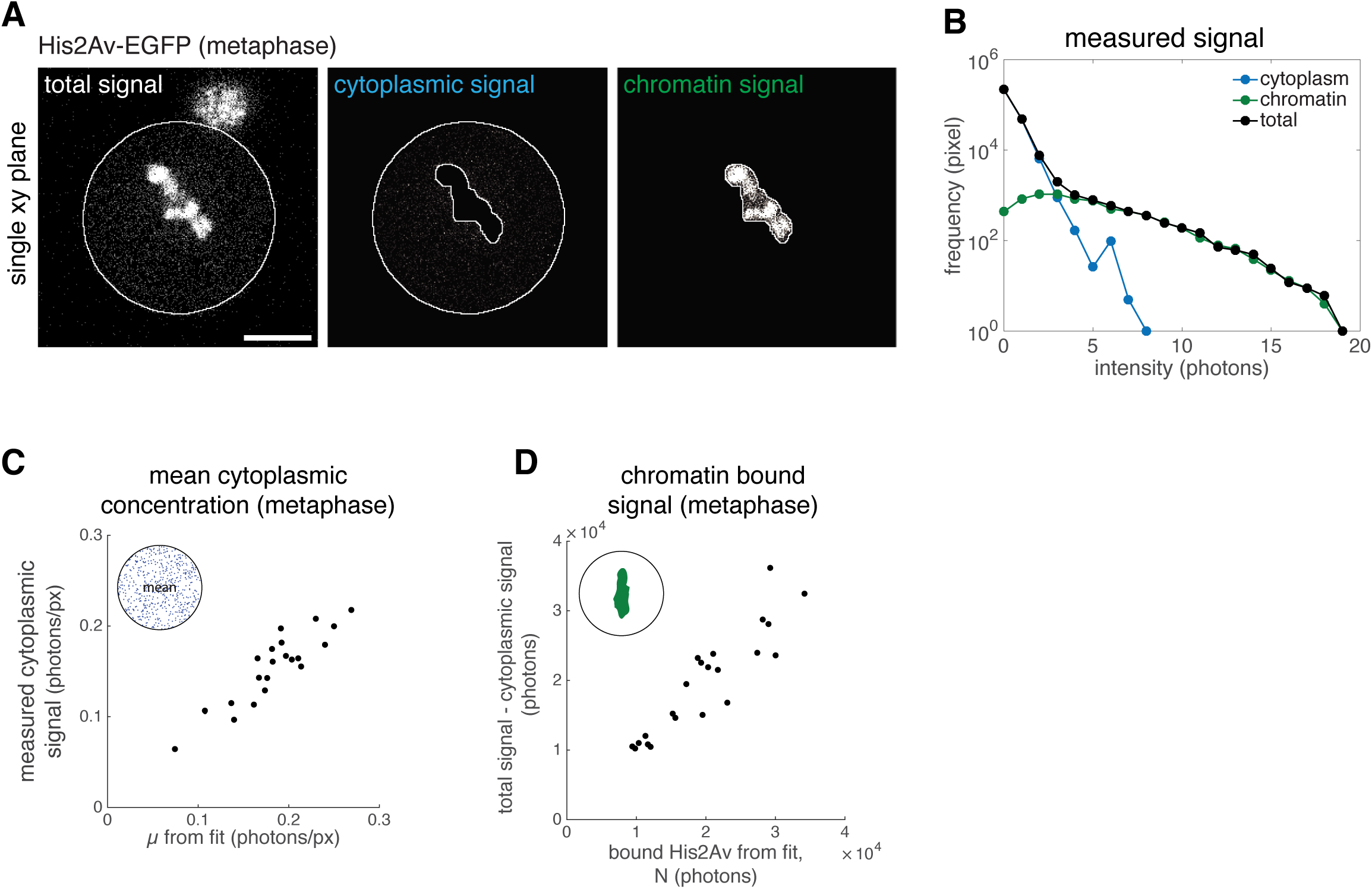
Fitting procedure and controls. **A)** Representative single planes of the cell in Figure 3A with manual segmenting of the cytoplasm as well as the chromatin. **B)** Histograms of total, cytoplasmic and chromatin intensities of the entire cell corresponding to the Fig S3-A. **C)** Mean cytoplasmic His2Av-EGFP signal during metaphase versus mean *μ* obtained from fit of representative cells of different sizes. We observe a linear correlation validating our fitting procedure. **D)** Measured chromatin signal (total intensity – cytoplasmic intensity) at metaphase versus bound His2Av signal after NEB obtained from fitting parameters (p,r). We observe a linear correlation validating our fitting procedure.

**Figure S4:**
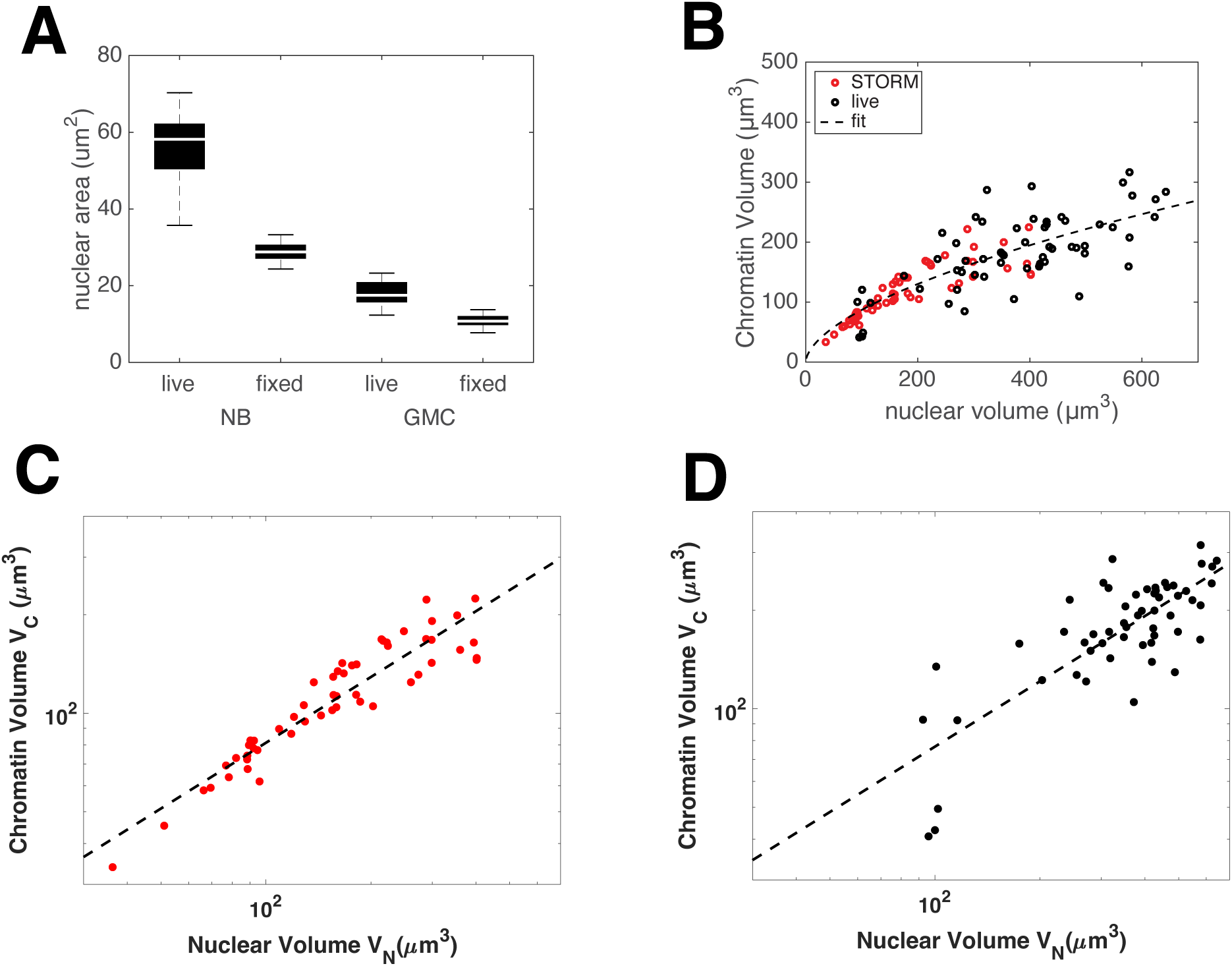
**A)** Comparison of live and fixed NBs and GMCs nuclear areas in interphase. A factor 2 is observed probably emerging from the fixation treatment. **B)** Graph showing the chromatin volume as a function of the nuclear volume in interphase in fixed (STORM) and live imaged cells. A power law fit on the combined dataset is displayed. The figure corresponds to the Figure 4A **C)** A power law fit on calculated interphase chromatin volume from fixed cells (STORM). The fitting equation is: 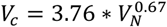 **D)** A power law fit on interphase chromatin volume from live imaging experiments. The fitting equation is: 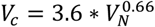

**Movie 1:** Time series of sum intensity projections of a cell cluster showing the division of chromosomes in an asymmetric division of a neuroblast expressing His2Av-EGFP

**Movie 2:** Time series of sum intensity projections of a neuroblast expressing His2Av-EGFP treated with 1 µM colcemid undergoing mitosis.

